# Age-related decrease in appetitive associative memory in fruit flies

**DOI:** 10.1101/2022.08.23.504945

**Authors:** Christian König, Bertram Gerber

## Abstract

Memory scores are dynamic across developmental stages. In particular, memory scores typically decrease from late adolescence into old age, reflecting complex changes in mnemonic and sensory-motor faculties, metabolic and motivational changes, and changes in cognitive strategy as well. In *Drosophila melanogaster*, such age-related decreases in memory scores have been studied intensely for the association of odours with electric shock punishment. We report that odour-sucrose reward memory scores likewise decrease as the flies age. This was observed after one-trial and after two-trial conditioning, and for both immediate testing and for recall tests one day later. This decrease was already particularly pronounced in relatively young animals, in the first 2-3 weeks after adult hatching, and was more pronounced in female than in male flies.

## INTRODUCTION

In animals and humans alike, memory scores typically decrease from adolescence via adulthood and old age into senescence, reflecting a combination of a wide variety of processes (reviewed in Bishop et al. 2010, Grady 2012). These include degrading mnemonic ability and sensory-motor faculties, motivational and metabolic changes, as well as changes in cognitive strategy. Regardless of the relative contribution of these processes, the age-related decrease in memory scores is observed not only across species, but across experimental paradigms, sensory modalities, valence and temporal domains, and in both sexes. It was therefore surprising to realize that for *Drosophila melanogaster*, a versatile study case for understanding ageing and age-related memory impairment (reviewed in He & Jasper 2014, Piper & Partridge 2018), most studies have focused on age-related decreases in aversive, odour-electric-shock associative memory, following the discovery of such decreases by Tamura et al. (2003) (for earlier reports in other, likewise aversively motivated paradigms: Fresquet & Medioni 1993, Guo et al. 1996, Savvateeva et al. 1999). In contrast, Tonoki et al. (2020) concluded that there is no age-related decrease in odour-sucrose associative memory in *Drosophila* (for details see Discussion section). Here, we report such a decrease, using a study design comprising three experimental conditions with five age cohorts each.

## MATERIALS AND METHODS

### Flies and fly husbandry

*Drosophila melanogaster* of the Canton Special wild-type strain were kept in mass culture on standard cornmeal-molasses food at 60 – 70% relative humidity and 25 °C temperature under a 12 h/ 12 h light/ dark cycle. Freshly hatched adults were collected in food vials and kept for the number of days indicated in the Results section. To prevent the flies getting stuck too frequently in the food once the food was churned up by their larval offspring, and to remove them long before their own offspring hatched as adults, the flies were placed on fresh food vials every 2-3 days.

### Behavioural experiments

The behavioural experiments followed standard methods as described in detail in Yarali & Gerber (2010), unless mentioned otherwise. In brief, 14-18 h before training started, mixed-sex cohorts of flies were transferred from their food vials to a vial without food but with only a moist tissue paper soaked with tap water (‘wet starvation’). Training and testing of the flies then took place using a custom-made apparatus (CON-ELEKTRONIK, Greussenheim, Germany) at 23 – 25 °C and 60 – 80% relative humidity. Training was performed in white room light, testing in red light invisible to the flies. As odorants, 50 µL benzaldehyde (BA) and 250 µL 3-octanol (OCT) (CAS 100-52-7, and 589-98-0, respectively; Merck, Darmstadt, Germany) were applied undiluted to 1-cm-deep Teflon containers of 5 and 14 mm diameter, respectively. From these, odour-loaded air could be shunted into the permanent airstream flowing through the apparatus.

Flies were trained and tested *en masse*, in cohorts of ∼100 individual animals each. For training, the flies were gently loaded into the setup. After one minute the flies were transferred to a training tube lined with a filter paper that had been soaked the day before with 2 ml of 2 M sucrose (D-sucrose; CAS: 57-50-1, Roth, Karlsruhe, Germany) solution and left to dry overnight. At the same time the trained odour (CS+) was shunted into the permanent airflow running through this tube. After 45 s, the odour presentation was terminated, and after an additional 15 s, the flies were removed from the sucrose-containing tube. At the end of a 1-min waiting period, the flies were transferred into another training tube, which was empty. At the same time, the reference odour (CS-) was presented. After 45 s, the odour presentation was finished, and 15 s later the flies were removed from this training tube. As indicated in the Results section, we used either one or two such training trials. The use of BA and OCT as CS+ and CS-was balanced across repetitions of the experiment, resulting in reciprocally trained groups of flies. In addition, both the sequence of the CS+/CS-presentations during training trials and the sequence of running the experiment for the reciprocal training groups were balanced, as described in König et al. (2019, *loc. cit. Supplemental Fig 1B*). To measure immediate memory, the flies were transferred after an additional waiting period of 3 min to a T maze, where they could choose between the trained (CS+) and the reference odour (CS-). After 2 min testing time, the arms of the maze were closed and the flies on each side were counted to calculate a preference index (BA PREF):

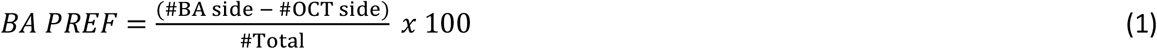

Thus, positive scores indicate a relative preference for BA and negative scores a relative preference for OCT. From the BA PREF scores of reciprocally trained groups of flies, that is of flies for which either BA or OCT was paired with the sucrose reward, a memory score was calculated as follows:

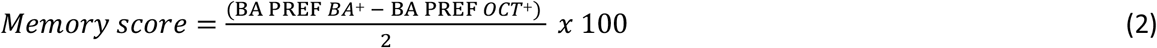

Positive values for the memory score thus reflect appetitive associative memory, negative values aversive associative memory.

As indicated in the Results section, in some cases the flies were tested the next day. In order to ensure reasonable survival rates, in these cases the flies were removed from the training apparatus and first kept for 1 h on standard food; only then were they wet-starved until the test took place the following day.

### Statistical analyses

Two-tailed non-parametric statistics were used throughout. To compare across multiple independent groups, Kruskal-Wallis tests (*H* tests) with subsequent pair-wise comparisons, Mann-Whitney *U*-tests (*U*-tests), were used (Statistica 13, RRID: SCR_014213, StatSoft Inc, Tulsa, USA). For comparisons of data from a group to chance levels (zero), one-sample sign tests (OSS) were applied. To keep the within-experiment error rate below 5 %, a Bonferroni-Holm correction for multiple comparisons was employed. The experimenter was blind to experimental groups during the counting of flies after the experiments. The data are presented as box plots showing the median as the middle line, the 25 and 75 % quantiles as box boundaries, and the 10 and 90 % quantiles as whiskers. An account of the data is provided in the Supplemental Data file ‘*Table S1 -Raw data*.*xlsx*’.

## RESULTS AND DISCUSSION

Flies in 6-day age cohorts from the day of adult hatching to 30 days after hatching were starved for 14-18 h and received a single trial of differential conditioning such that one odour was paired with a sucrose reward, whereas the other odour was not. Then the flies were tested for their relative preference between the two odours. From these preferences, associative memory scores were calculated that reflect their learned preference for the previously reward-associated odour (Figure 1A) (using a reciprocal design, the chemical identity of the odours was balanced across repetitions of the experiment). As shown in Figure 1B, memory scores decreased across age cohorts: relative to the youngest cohort, memory scores were reduced from an age of 13-18 days on (see Supplemental Figure 1A for underlying preference scores). The results were similar when two conditioning trials were used (Figure 1C and Supplemental Figure 1B) (in this case a decrease was already significant for the 7-12 day age cohort) and also when after such two-trial differential conditioning 24 hours were allowed to pass until testing (Figure 1D and Supplemental Figure 1C). Notably, memory scores remained significantly above chance levels, i.e. were larger than zero, for all cases with the exception of the 24-hour memory scores in the very oldest age cohort.

**Fig. 1:**
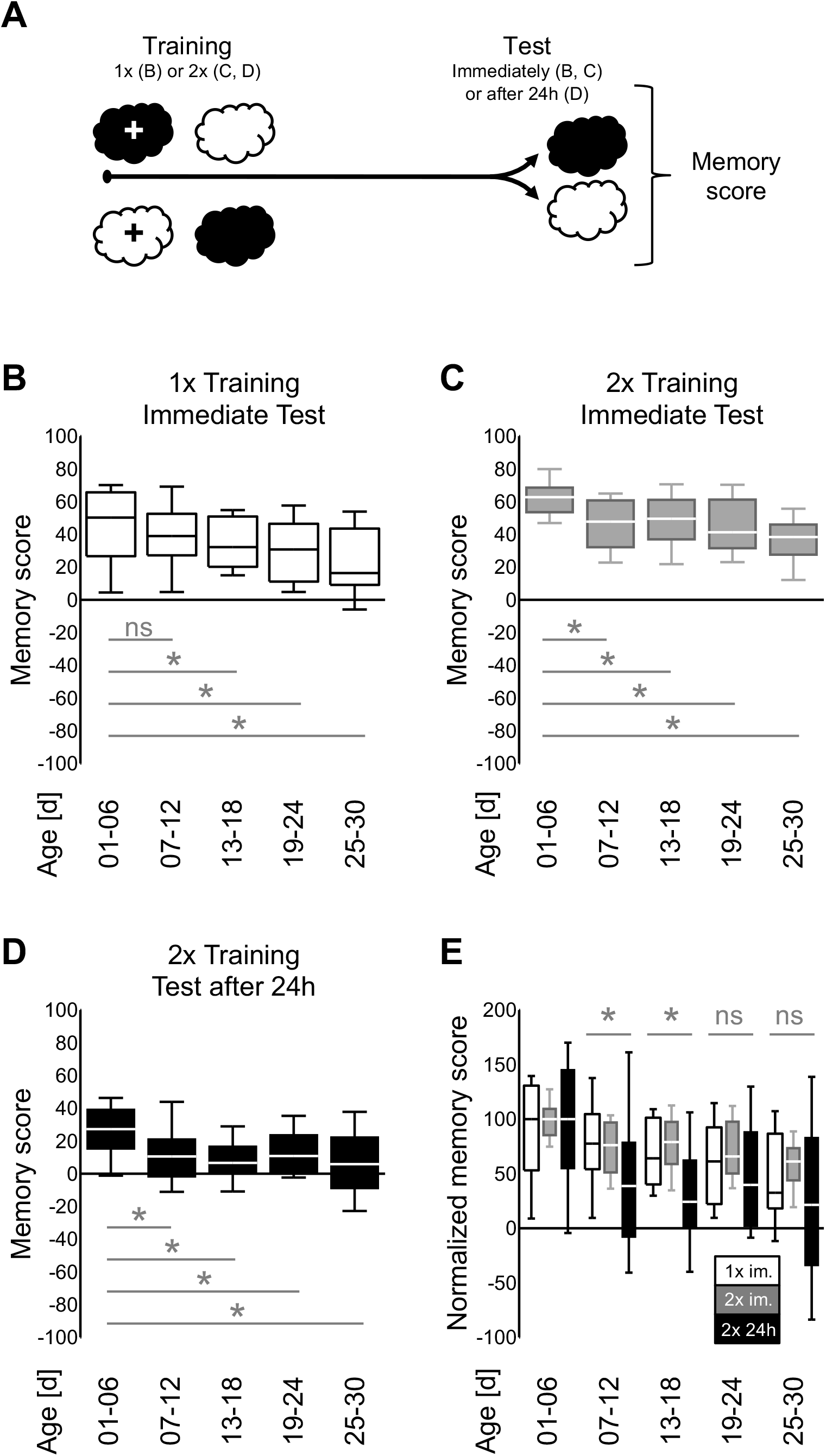
Age-related decrease in appetitive associative memory scores. **A)** Flies of the age cohorts indicated in (B-E) were differentially conditioned such that during training one but not the other of two odours (white and black clouds) was presented with a sucrose reward (+); the experiment followed a reciprocal design, meaning that the chemical identity of the odours was swapped across repetitions of the experiment. During the test, the flies could choose between the two odours in a T-maze, and their preference between the two odours was noted. Associative memory is reflected in the difference in preference between reciprocally trained sets of flies (quantified through the memory scores). By convention, positive and negative memory scores in (B-E) indicate appetitive and aversive associative memory, respectively. Please note that the sequence of events during training was as indicated in half of the cases (i.e. the initially presented odour was paired with the sucrose, whereas the odour presented second was not); this was reversed in the other half of the cases. **B)** After one training trial and when tested immediately after training, memory scores differed across age cohorts (N= 43, 39, 32, 39, 27). Relative to the youngest age cohort of 1-6 days after adult hatching, the memory scores were reduced for the age cohorts of 13-18 days or older. **C)** After two training trials too, memory scores immediately after training differed across age cohorts (N= 46, 37, 41, 36, 34). Relative to the youngest age cohort, the memory scores in this case were reduced for all older age cohorts. **D)** As in (C), but the testing was 24 hours after training. Memory scores differed across age cohorts (N= 38, 39, 59, 40, 64); relative to the youngest age cohort the memory scores were reduced in all cases. **E)** Data from (B-D) normalized to the median memory score of the youngest age cohort of the respective experimental condition. Normalized memory scores differed between experimental conditions for the 7-12 and the 13-18 day age cohort, but not for the older age cohorts. In (B-D), * refers to *P*< 0.05 and ns to *P*> 0.05 in *U*-tests of the youngest age cohort versus the older age cohort indicated, corrected for multiple testing according to Bonferroni-Holm; all memory scores except those for the oldest age cohort in (D) were significantly different from chance levels, i.e. from zero, in OSS-tests (*P*< 0.05, corrected according to Bonferroni-Holm). In (E), * refers to *P*< 0.05 and ns to *P*> 0.05 in KW-tests across the three experimental conditions for the age cohort indicated, corrected for multiple testing according to Bonferroni-Holm. Box plots represent the median as the middle line, 25 and 75% quantiles as box boundaries, and 10 and 90% quantiles as whiskers. Preference scores underlying the memory scores are documented in Supplemental Figure 1. All statistical results and the data are documented in *Table S1 -Raw data*.*xlsx*.

As expected, the memory scores for the youngest age cohort differed across the three experimental conditions (1-trial memory, 2-trial memory, 2-trial^24h^ memory; KKW test: H(2, N=127)=45.62, P<0.05; MWU tests: 1-trial im. vs. 2-trial im. U=630, P<0.05; 1-trial im. vs. 2-trial^24h^ U=431, P<0.05/2; 2-trial im. vs. 2-trial^24h^ U=114, P<0.05/3) (Schwaerzel et al. 2003, Kim et al. 2007, Colomb et al. 2009, Tonoki et al. 2020). In order to compare the rate of the age-related decrease in memory scores across these three experimental conditions, we therefore normalized all the memory scores for each of the three experimental conditions to the median of its respective youngest age cohort. This showed that the age-related decrease in memory scores was strongest for 2-trial^24h^ memory, in particular across days 7-18 (Figure 1E). Of note is that the more partial decrease in memory scores for 1-trial and 2-trial memory in this 7-18 day age-window was more pronounced in females than in males (Figure 2, Supplemental Figure 2) (for the even older age cohorts a separation by sex was not carried out, because the number of surviving flies, in particular the number of males, appeared to be too low, leading to excessively variable memory scores).

**Fig. 2:**
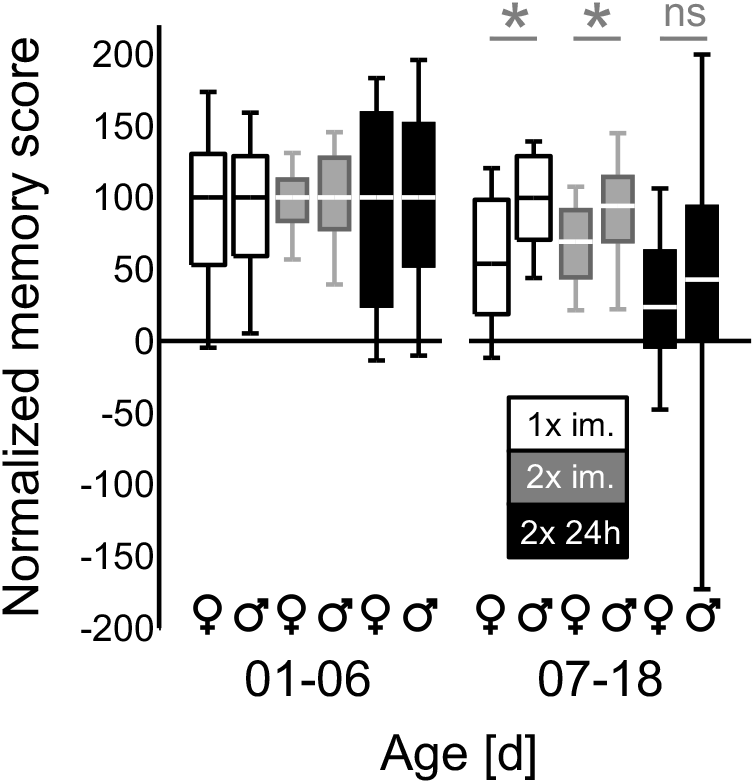
The age-related decrease in memory scores is more pronounced in females than in males. The data from Figure 1E are presented separated by sex, for the youngest age cohort and for the combined age window of 7-18 days after adult hatching. The data were normalized, separately for each sex, to the median memory score of the 1-6 day age cohort of the respective experimental condition. For 1-trial and 2-trial memory, but not for 2-trial^24h^ memory, this revealed that memory scores decreased more strongly in female than in male flies (N= 43, 43, 46, 46, 38, 38, 71, 71, 78, 78, 96, 79) * refers to *P*< 0.05 and ns to *P*> 0.05 in *U*-tests between the sexes, corrected for multiple testing according to Bonferroni-Holm. Other details as in Figure 1. All statistical results and the data are documented in *Table S1 -Raw data*.*xlsx*.

Thus, across experimental conditions the present study reveals an age-related decrease in appetitive associative memory scores during the first 2-3 weeks after adult hatching (Figure 1E), which is more pronounced in females than in males (Figure 2). While the present study does not attempt to disentangle what are arguably multiple chains of cause and effect leading to this decrease, it suggests that future research could focus on this relatively early time window when looking into the interplay of sensory, motor, motivational, metabolic, mnemonic and cognitive processes contributing to age-related decreases in appetitive memory scores. For the practitioner, such an early time window of the effect is good news because the experiments take less time than those focusing on later stages, and require less attention to fly husbandry. Also, the contribution of non-target age-related effects is conceivably more pronounced for older stages. On the other hand, decreases in memory scores at this relatively young age might be of limited translational validity for processes at an older age that are arguably more relevant from a biomedical point of view.

In contrast to the present study, Tonoki et al. (2020) concluded that there is no age-related decrease in memory scores for the association of odours with sucrose in *Drosophila*. Although their experimental paradigm is in principle similar to the one used here, there are a number of procedural differences. These include aspects of the starvation procedure and of odour presentation, of the light conditions during training, of the duration of the feeding period between training and test for the measurement of 24-hour memory, and of the sequence of CS+/CS-presentations during training. These sequences were balanced in the present study but always started with CS-in Tonoki et al. (2020). For the relatively young age groups of 2 versus 10 days of age, moreover, Tonoki et al. (2020, *loc. cit.* Fig. 1E-G) used a more extended starvation period of 40 hours as compared to the 14-18 hours used in the present study, which did uncover a decrease in memory scores in flies over roughly that age range (Figure 1E). Strikingly, using a shorter starvation period of 18 hours, similar to the 14-18 hours used here, Tonoki et al. (2020) found higher (*sic*) memory scores for 30-day-old than for 10-day-old flies (*loc. cit. Fig. 1C*) (compare to Figure 1E). Which of the above-mentioned differences in procedure might account for the discrepancies in results must remain unresolved for now. Concerning the more pronounced age-related decrease in memory scores in females than in males that we report (Figure 2), it seems significant that females are less susceptible to starvation than males (Robinson et al. 2000). Accordingly, for an older age at least (7-18 days), our data suggest that survival rates for the 24-hour memory test are better in female than in male flies (Supplemental Figure 2). Thus, at that age the nutritious sucrose reward received during training might be less motivationally salient, and indeed less rewarding, to the females than to the males.

Given that the decrease in memory scores uncovered here both takes place relatively early in adult life (Figure 1E) and is more pronounced in females than in males (Figure 2), one wonders whether female life-history traits play a role. Indeed, Scheunemann et al. (2019) have shown that memory scores after aversive associative learning between odours and electric shock are more negative for mated than for non-mated females. As the proportion of mated females increases across early adult life, it seems possible that the less positive memory scores observed here for 7-18-day-old than for younger female flies may partially reflect the fact that already mated, older females establish a relatively more negative ‘take home message’ both from the appetitive training procedure employed here and from the aversive training procedure of Scheunemann et al. (2019).

## ACKNOWLEDGEMENTS

We thank Bettina Kracht and Anna Ciuraszkiewicz for technical assistance, Bert Klagges, Thomas Niewalda and Naoko Toshima for comments on earlier versions of the manuscript, and R. D. V. Glasgow for language editing.

## AUTHOR CONTRIBUTIONS

BG: Conceptualization, Validation, Writing – review editing, Project administration, Visualization, Funding acquisition

CK: Conceptualization, Validation,, Formal analysis, Data curation, Visualization, Writing – original draft

## Supplemental Figures

**Figure S1.**
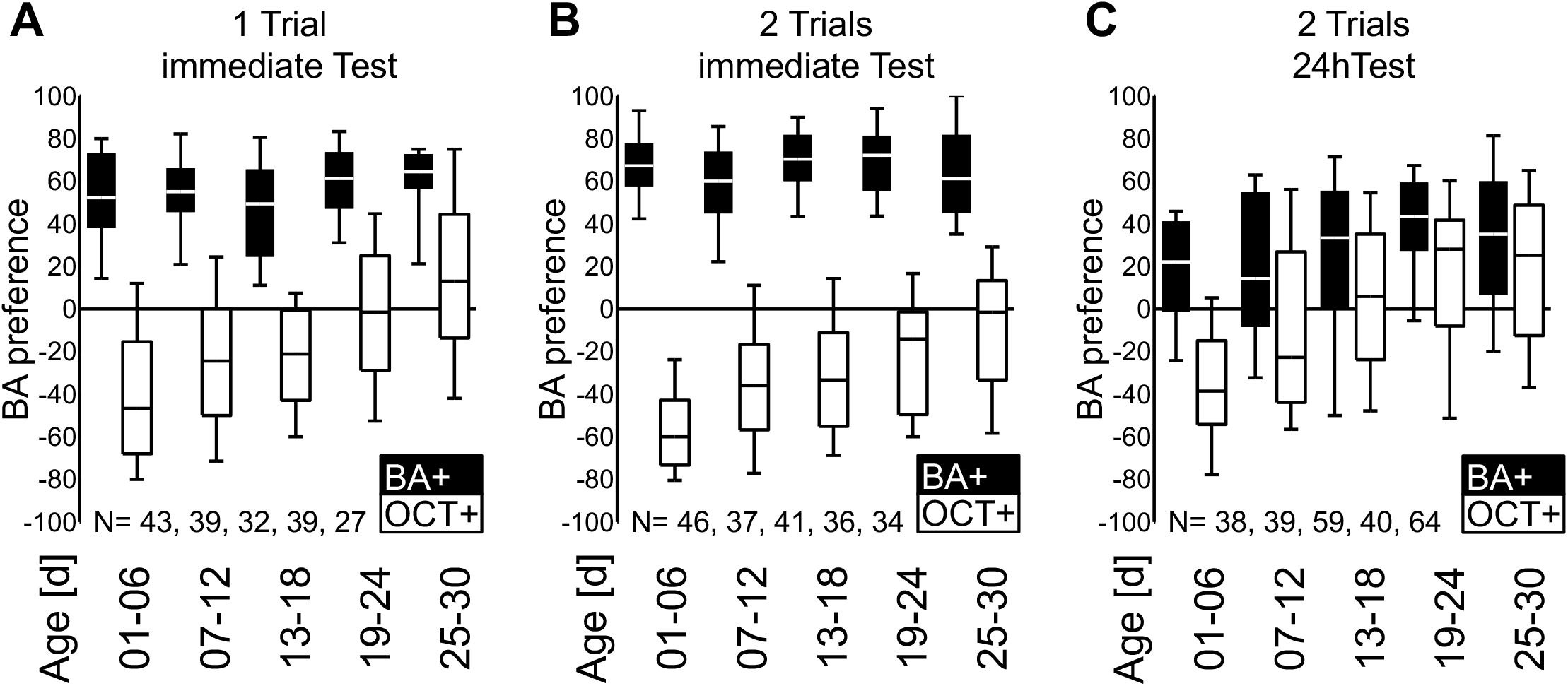
BA preferences underlying the memory scores in Figure 1B-D. Panels (A-C) show BA preference data underlying the memory scores shown in Fig. 1B-D, respectively. Filled boxes represent BA preference in a BA versus OCT choice test after either BA or OCT was paired with the sucrose reward during training (BA+ or OCT+, respectively). Box plots represent the median as the midline, 25 and 75% as the box boundaries, and 10 and 90% as the whiskers. Sample sizes are indicated within the figures. The data are documented in Table S1 -Raw data.xlsx.

**Figure S2.**
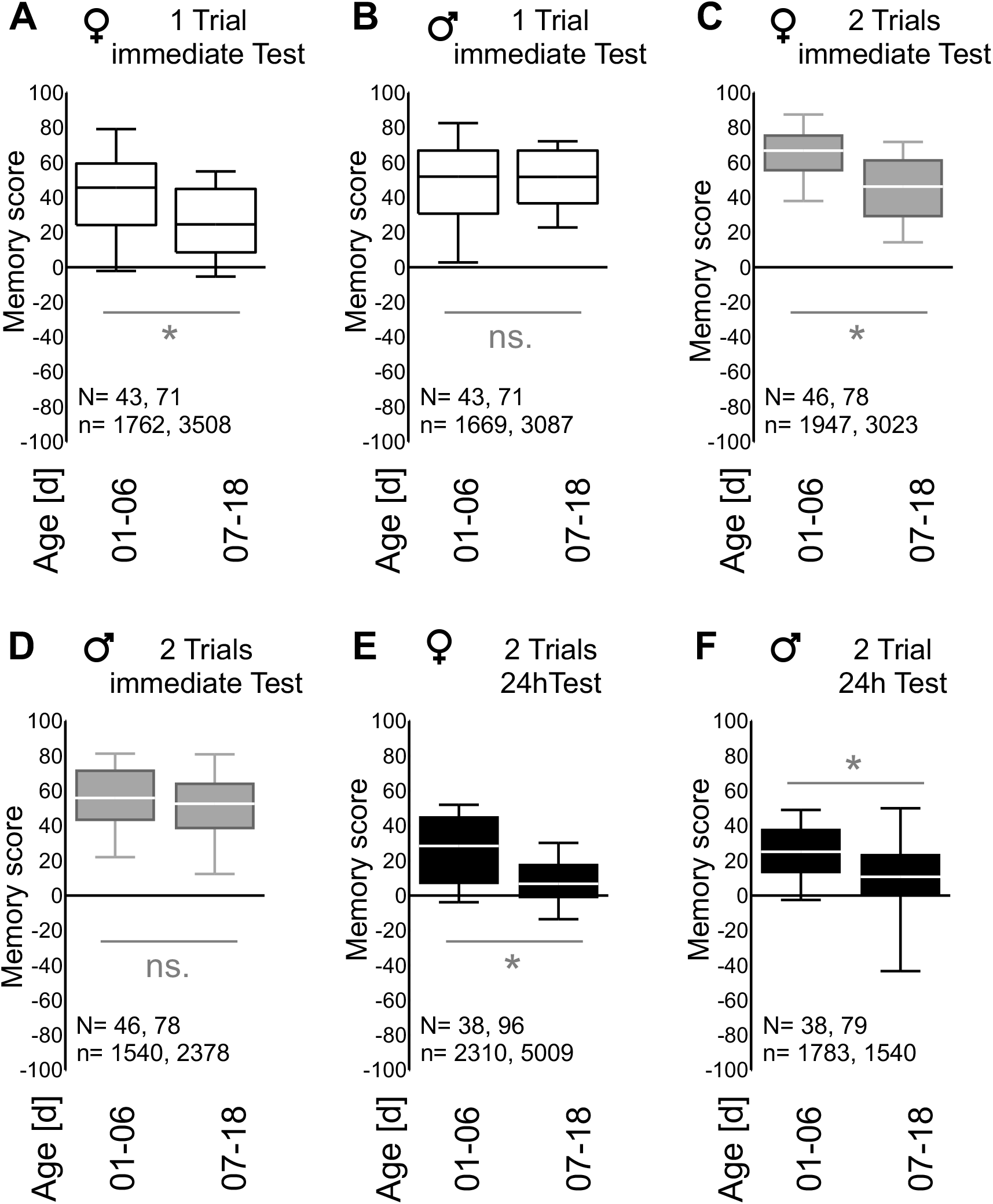
Memory scores underlying the normalized memory scores in Figure 2. Shown are the memory scores, separated by sex, that are the basis for the normalized memory scores shown in Fig. 2. (A-D) For memory after 1-trial (A and B) and 2-trial training (C and D) only female but not male memory scores are reduced in the older age cohort. (E and F) For 2-trial^24h^ memory both female and male memory scores are reduced in the older age cohort. Sample sizes (N) and number of total individual flies (n) are indicated within the figures. Note the relatively lower number of surviving male flies per sample for the 24-hour memory test (E, F) (on average 52 female flies per sample as compared to on average 19 male flies per sample; in 17/96 samples indeed *only* female flies had survived). * refers to P< 0.05 and ns to P> 0.05 in U-tests between the age cohorts, corrected for multiple testing according to Bonferroni-Holm. All memory scores were significantly different from chance levels, i.e. from zero, in OSS-tests (P< 0.05, corrected according to Bonferroni-Holm). Other details as in Figure S1. All statistical results and the data are documented in Table S1 -Raw data.xlsx.

